# SOX17 Is Not Required for the Derivation and Maintenance of Mouse Extraembryonic Endoderm Stem Cell Lines

**DOI:** 10.1101/2022.01.09.475525

**Authors:** Xudong Dong, Ailing Ding, Jiangwei Lin

## Abstract

Extraembryonic endoderm stem (XEN) cell lines can be derived and maintained in vitro and reflect the primitive endoderm cell lineage. SOX17 is thought to be required for the derivation and maintenance of mouse XEN cell lines. Here we have re-evaluated this requirement for SOX17. We derived multiple SOX17-deficient XEN cell lines from preimplantation embryos of a SOX17-Cre knockout strain and chemically converted multiple SOX17-deficient embryonic stem cell lines into XEN cell lines by transient culturing with retinoic acid and Activin A. We confirmed the XEN profile of SOX17-deficient cell lines by immunofluorescence with various markers, by NanoString gene expression analyses, and by their contribution to the extraembryonic endoderm of chimeric embryos produced by injecting these cells into blastocysts. Thus, SOX17 is not required for the derivation and maintenance of XEN cell lines.

## INTRODUCTION

During mouse embryogenesis, the morula undergoes compaction and gradually differentiates into the trophectoderm (TE) around the outer cell mass and the inner cell mass (ICM) (Johnson and Ziomek, 1981). By embryonic day E3.5, the second cell fate decision takes place involving the segregation of ICM into the pluripotent epiblast (EPI) and primitive endoderm (PrE), which are distributed in a salt-and-pepper pattern. By the late blastocyst stage, PrE forms a layer of cells along the surface of the blastocoel cavity. The epiblast gives rise to the embryo proper, amnion, and extraembryonic mesoderm of the yolk sac. TE cells give rise to the placenta. PrE forms the two ExEn lineages: visceral endoderm (VE) and parietal endoderm (PE) of the yolk sac (Chazaud et al., 2006; Plusa et al., 2008; Artus and Hadjantonakis, 2012). VE cells and their derivatives play a critical role in organogenesis. VE cells are the first site of hematopoiesis (Toles et al., 1989; McGrath and Pails 2005) and form blood islands and endothelial cells through the expression of Indian hedgehog and vascular endothelial growth factor (VEGF) (Byrd et al., 2002; Damert et al., 2002). At gastrulation, VE cells contribute to forming the gut endoderm tissue of the fetus (Kwon et al., 2008). VE and PE function as an “early placenta” that is responsible for nutrient and waste exchange (Cross et al., 1994).

Extraembryonic endoderm stem (XEN) cell lines can be derived and maintained in vitro (Kunath et al., 2005; Niakan et al., 2013) and reflect the PrE cell lineage. There are four methods to derive mouse XEN cell lines. First, XEN cell lines can be derived directly from blastocysts (Kunath et al., 2005). Second, XEN cell lines can be converted from embryonic stem (ES) cells by forced expression of XEN-specific genes such as *Gata6* (Wamaitha et al., 2015), *Gata4* (Fujikura et al., 2002), or *Sox17* (McDonald et al., 2014) or chemically by transient culturing with retinoic acid (RA) and activin A (Cho et al., 2012). Third, XEN cell lines can be induced from fibroblasts by overexpression of classical OSKM factors (Parenti et al., 2016). Fourth, we have reported the efficient derivation of XEN cell lines from postimplantation embryos (Lin et al., 2016, 2017).

SOX17 is a member of the Sry-related high-mobility group box (Sox) transcription factors and has an essential role in the differentiation of several types of cells (Foster et al., 1994). During mouse embryogenesis, SOX17 is first detected at the 16–32 cell stages coexpressed with Oct4, then in the PrE of blastocysts, and later in VE at E6.0 and in the endoderm at E7.5, where it plays an essential role in organ formation (Kanai-Azuma et al., 2002). Previous studies have revealed its role in the regulation of fetal hematopoiesis (Kim et al., 2007) and vasculogenesis (Matsui et al., 2006; Sakamoto et al., 2007). SOX17 has also been proposed to function as a key regulator of endoderm formation and differentiation, a function that is conserved across vertebrates (Hudson et al., 1997; Alexander et al., 1999; Clements et al., 2000). In mice, genetic inactivation of SOX17 leads to severe defects in the formation of the definitive endoderm (Kanai-Azuma et al., 2002). Overexpression of SOX17 is sufficient to promote ES cells to convert to XEN cells (McDonald et al., 2014). SOX17 is critical for PrE formation, and a lack of SOX17 will significantly decreases the PrE numbers of blastocysts (Artus et al., 2011). XEN cell lines cannot be derived from SOX17 mutant embryos and converted from ES cells (Niakan et al., 2010; Cho et al., 2012). Downregulation of SOX17 by RNA interference will impairs XEN cell maintenance (Lim et al., 2008). Embryonic bodies derived from SOX17 mutant ES cells fail to correctly form the outer ExEn layer (Niakan et al., 2010). SOX17 mutant ES cells differentiate into PrE cells but fail to differentiate into PE and VE fates (Shimoda et al., 2007). Here, we re-evaluated the requirement for SOX17 in the derivation and maintenance of XEN cell lines.

The model of sequential expression of PrE lineage-specific genes is *Gata6 > Pdgfra > Sox17 > Gata4 > Sox7* (Artus et al., 2010, 2011). Cells that express SOX17 can be visualized in a gene-targeted knockout mouse strain in which a fusion protein with green fluorescent protein (GFP) and Cre is expressed from the SOX17 locus (Choi et al., 2012) (although this strain is reported to contain and express GFP, and we confirmed the presence of GFP in this targeted insertion in the SOX17 locus, but we cannot detect GFP expression in embryos and cell lines derived from them). In this strain, which we refer to as SOX17-Cre, while SOX17-Cre crosses the gene-targeted Cre reporter strain R26-tauGFP41 (Wen et al., 2011), the GFP reporter is coexpressed with endogenous SOX17 protein and with PrE markers GATA6, GATA4, and DAB2 in preimplantation embryos (Choi et al., 2012; Lin et al., 2016). GFP colocalizes in the same cells with PrE markers GATA6 and GATA4 in blastocysts cultured in vitro and is expressed in the visceral and parietal endoderm of postimplantation embryos (Choi et al., 2012; Lin et al., 2021). XEN cell lines derived from R26-tauGFP41 × SOX17-Cre heterozygous blastocysts display the intrinsic fluorescence of GFP (Lin et al., 2016, 2021). Thus, in this strain, GFP serves as a robust live marker for PrE and its extraembryonic endoderm derivatives and can be applied in the context of XEN cell line derivation.

## RESULTS

### Derivation of XEN Cell Lines from SOX17-Deficient Blastocysts

In our previous work (Lin et al., 2017), we obtained a PDGFRA-deficient XEN cell line from blastocyst ICMs. In the experiments, we collected 27 E1.5–E2.5 embryos from SOX17-Cre heterozygous female intercrosses with SOX17-Cre heterozygous male and cultured them in KSOM medium to the blastocyst stage. We then removed the zona pellucida using acid Tyrode solution. By immunosurgery (Lin et al., 2011), we isolated 27 ICMs from 27 blastocysts (Fig. 1A). We transferred each of 27 ICMs into a well of a 4-well dish, coated them with 0.1% gelatin, covered them with MEF, and cultured them in ES medium with LIF. Outgrowths formed on day 4 (Fig. 1B-C). We changed the medium every 2-3 days without passaging the cells. On day 7 the outgrowths got larger, and the ICMs in Fig. 1B showed that all most cells became XEN-like cells and in Fig. 1C showed two distinct cell types (the XEN-like cells were in the top of the outgrowth as pointed with in red arrow, and the ES-like cells were in the bottom of the outgrowth as marked with in blue arrow). We replaced the ES medium with LIF by TS medium with F4H (25 ng/ml FGF4 and 1 *μ*g/ml heparin) on day 7. On day 14, we disaggregated the outgrowths and passaged the cells into a well of a 4-well dish, coated them with gelatin and covered them with MEF. On day 21, the ICMs in Fig. 1B established XEN cell lines; however, in Fig. 1C, only a small number of XEN-like cells were observed in the culture (day 21). We picked XEN-like colonies on day 30 and passaged them to new plates, and then on day 50, we established the XEN cell line. Our strategy was to isolate ICMs and make the ICMs to form a large outgrowth and then disaggregate and passage the outgrowths on day 14. We reasoned that epiblasts of ICMs would convert to PrE in large outgrowths and that fewer ES-like cells would be present (Lin et al., 2021). We thus derived 21 XEN cell lines from 27 ICMs. The genotype results showed that Fig. 1B is SOX17-Cre heterozygous and that Fig. 1C is SOX17-Cre homozygous. Thus, we obtained 14 XEN cell lines that were genotyped as SOX17-Cre heterozygous, 5 XEN cell lines as wild type, and two XEN cell lines as SOX17-Cre homozygous (X-ICM-12 and X-ICM-16) (Fig. 1D). Immunofluorescence analysis indicated that X-ICM-12 was positive for the XEN cell markers DAB2, GATA4, GATA6, and SOX17 but negative for the ES cell markers OCT4 and NANOG (Figure 1E). The SOX17-Cre-deficient cell line X-ICM-12 (Fig. 1F) was immunonegative for SOX17, and the SOX17-Cre heterozygous cell line X-ICM-17 was immunoreactive for SOX17 (Fig. 1G).

**Figure 1.**
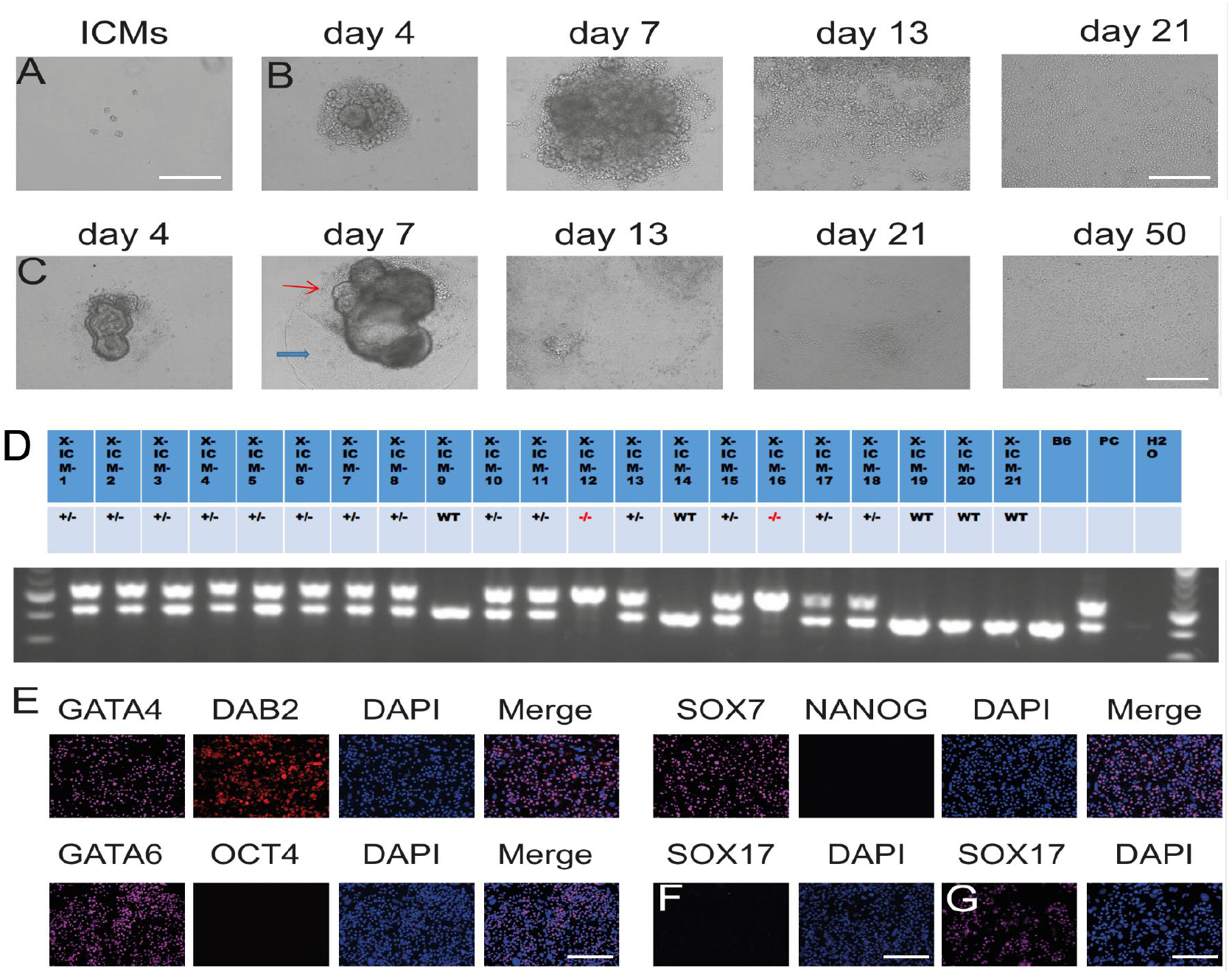
XEN Cell Lines Derived From SOX17-Deficient Blastocysts. (A) ICMs isolated from blastocyst. (B) XEN cell line X-ICM-17 derived from a SOX17-Cre heterozygous ICM. (C) XEN cell line X-ICM-12 derived from a SOX17-deficient ICM. (D) Genotyping results. Positive control: genomic DNA from the tail of a SOX17-Cre heterozygous mouse. Red: SOX17-Cre homozygous cell lines. (E) X-ICM-12. Immunofluorescence for GATA4, GATA6, SOX7, DAB2, OCT4, and NANOG. Third column: DAPI nuclear stain (blue). Fourth column: Merge. (F, G) X-ICM-12 is immunonegative (F), and X-ICM-17 is immunoreactive for SOX17 (G). Scale bars: 400 *μ*m in all panels.

### Conversion of SOX17-Deficient ES Cell Lines into cXEN cells

We set up SOX17-Cre heterozygous female intercrosses with SOX17-Cre heterozygous male. Using ES medium with LIF and 2i (PD0325901 and CHIR99021), we derived 4 SOX17-Cre heterozygous ES cell lines (ES-1, ES-3, ES-4, ES-7), 2 wild-type ES cell lines (ES-2, and ES-5) and 2 SOX17-Cre homozygous ES cell lines (ES-6 and ES-8) from 14 blastocysts. We obtained (R26-tauGFP41 x SOX17-Cre) F1 mice by establishing a natural mating between a homozygous female of the gene-targeted Cre reporter strain R26-tauGFP41 (Wen et al., 2011) and a heterozygous male of the gene-targeted driver strain SOX17-Cre. We set up a (R26-tauGFP41 x SOX17-Cre) F1 heterozygous intercross with a (R26-tauGFP41 x SOX17-Cre) F1 heterozygous. In (R26-tauGFP41 x SOX17-Cre) F2 embryos, cells would become permanently labeled with GFP upon expression of *SOX17* (which occurs in PrE but not in epiblasts), and the descendants of these cells, including XEN cells, would also be labeled permanently (Lin et al., 2021). Using ES medium with LIF and 2i, we derived 7 SOX17-Cre heterozygous ES cell lines (ES-9, ES-10, ES-11, ES-12, ES-14, ES-17, and ES-18), 1 wild-type ES cell line (ES-12) and 2 SOX17-Cre homozygous ES cell lines (ES-13 and ES-15) from 13 blastocysts.

We previously noticed that in PDGRFA-GFP ES cell lines, sparse GFP+ cells surrounded rare ES cell colonies (ES cells typically do not express PDGRFA and are thus GFP-) (Lin et al., 2017). The occurrence of these cells is in agreement with observations that SOX17 is expressed in a subset of cells on the outside of otherwise undifferentiated ES cell colonies (Niakan et al., 2010), that ES cells cultured in LIF and 2i contain a few cells expressing GATA6 (Morgani et al., 2013), and that PDGFRA-GFP heterozygous and homozygous ES cells contain a fraction of GFP+ cells (Lo Nigro et al., 2017). It thus appears that some ES cells convert spontaneously to XEN- or XEN-like cells.

A low dose of retinoic acid (RA) and Activin A promotes the chemical conversion of ES cells into XEN cells (termed as cXEN cells) but fails to convert SOX17-deficient ES cells into cXEN cells (Cho et al., 2012; Niakan et al., 2010). We followed the cXEN conversion protocol of Cho et al. 2012 with a slight change (Lin et al., 2017). The protocol for the conversion of cXEN from ES cells is shown in Fig. 2A. We cultured ES-9, ES-12 and ES-17 (R26-tauGFP41×SOX17-Cre F2 heterozygous) and ES-13 and ES-15 (R26-tauGFP41×SOX17-Cre F2 homozygous) for 48 hr in standard TS cell medium with F4H, to which 0.01 *μ*M RA and 10 ng/ml Activin A were added. Thereafter, all cells were cultured in standard TS cell medium with F4H. XEN-like colonies accumulated on days 6 and 11 (Fig. 2B). We picked XEN-like colonies and passaged the cells into new plates. On day 20, XEN-like cells accumulated but still had a small fraction of ES-like colonies in the plates. Passaging of the mixed cells to single cells by trypsin releases ES-like colonies to many ES-like cells and grows many ES-like colonies, and then the ES-like cells dominate the population. In this case, we used a method that is called the crowded culture method or extended the culture method (Lin et al., 2017). We changed the medium every other day but passaged the cells every 1-2 weeks. As the cultures grew confluent, a fraction of the GFP+ (XEN-like) cells did not adhere tightly to the dishes and were easier to lose during medium changes. It appears that whereas colonies of ES-like cells and differentiating ES cells adhered tightly to the dishes, XEN-like cells became sorted to the outside of these colonies (Artus et al., 2012) and then were excluded from the colonies. We therefore systematically enriched GFP+ (XEN-like) cells when a medium change was due to spinning down the suspended cells and transferring them into a new dish coated with gelatin and covered with MEF. We thus converted cXEN cell lines from SOX17-Cre heterozygous ES cell lines (ES-9, ES-12 and ES-17) after ~21 days and from SOX17-deficient ES cell lines (ES-13 and ES-15) after ~30 days. The SOX17-deficient cXEN cell line that we converted from ES-13 (called cXEN-13) was maintained for >60 days (Fig. 2B), and the XEN cell phenotype kept in culture. Next, we applied the same protocol to convert SOX17-deficient ES cell lines ES-6 and ES-8 (SOX17-Cre homozygous) cells into cXEN cell lines. After ~30 days, we obtained a stable cXEN cell line from each ES cell line, called cXEN-6 and cXEN-8. Immunofluorescence analysis indicated that the four SOX17-deficient cXEN cell lines were positive for the XEN cell markers DAB2, GATA4, GATA6, and SOX7, but negative for the ES cell markers OCT4 and NANOG (Fig. 2C). The SOX17-deficient cell line cXEN-13 was immunonegative for SOX17 (Fig. 2D), and the SOX17-Cre heterozygous cell line cXEN-12 was immunoreactive for SOX17 (Fig. 2E).

**Figure 2.**
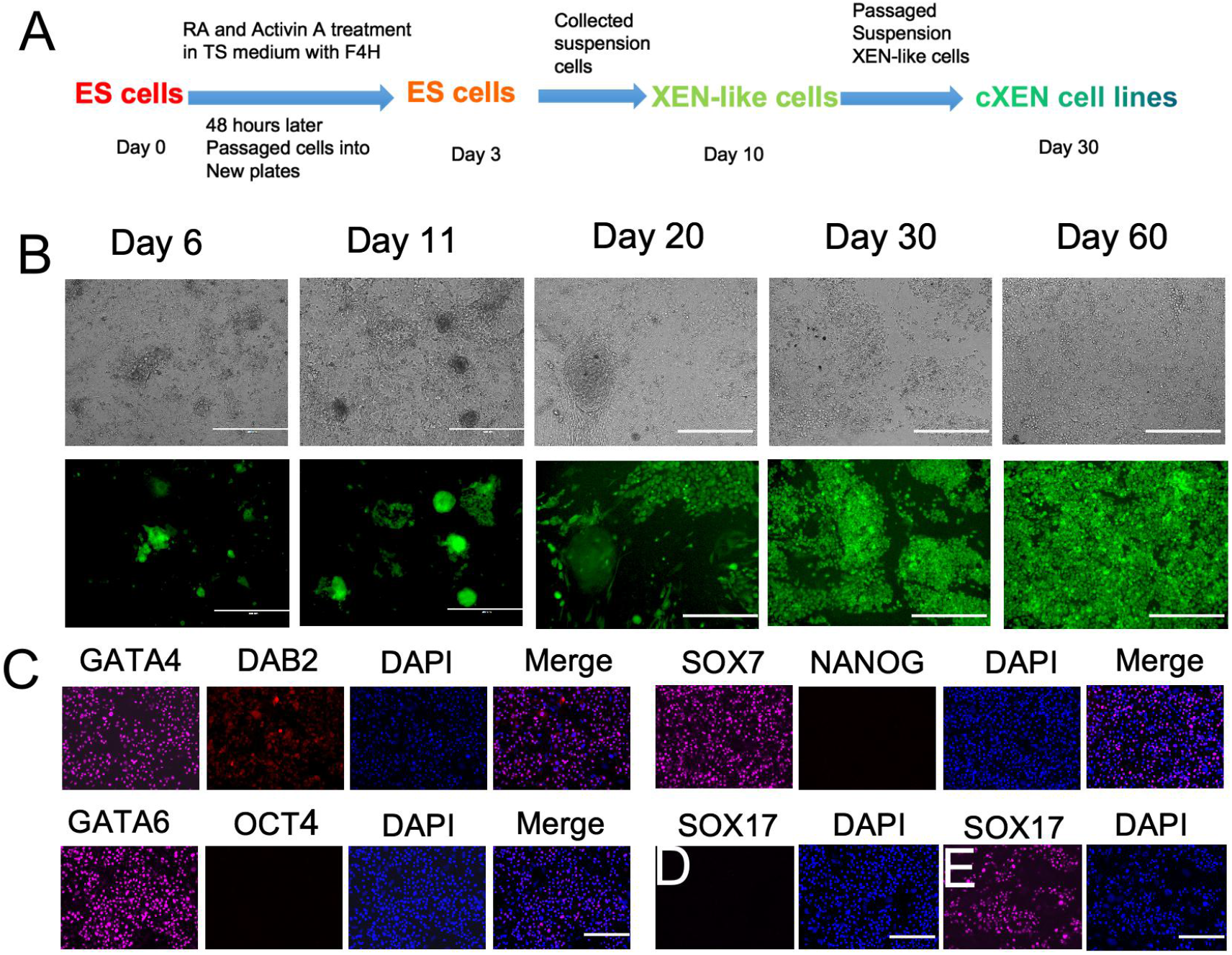
cXEN Cell Lines Converted Chemically From SOX17-Deficient ES Cell Lines. (A) Schematic representation showing that ES cells converted into the cXEN cell line treated with RA and Activin A. (B) Conversion of ES-13 (SOX17-deficient) into cXEN cells in TS medium with F4H, 0.01 *μ*M RA, and 10 ng/ml Activin A. (C) Immunofluorescence on SOX17-deficient cXEN-13 cells converted from ES-13. Immunofluorescence for GATA4, GATA6, SOX7, DAB2, OCT4 and NANOG. (D, E) cXEN-13 is immunonegative for SOX17 (D), and cXEN-12 is immunoreactive for SOX17 (E). Scale bars: 400 *μ*m in all panels.

### NanoString Gene Expression Analyses of XEN Cell Lines and ES Cell Lines

Next, we applied the NanoString multiplex platform (Khan et al., 2011, Lin et al. 2016, 2017) to compare patterns of gene expression in SOX17-Cre homozygous ES and XEN cell lines. All XEN cell lines had high levels of expression of XEN cell-specific genes, such as *Gata4, Gata6, Pdgfra, Sox7*, and *Dab2*, versus low levels of expression or no expression of ES cell-specific genes, such as *Sox2, Pou5f1* / *Oct4, Nanog*, and *Zfp42/Rex1* (Fig. 3). In SOX17-Cre homozygous XEN cell lines, *Sox17* expression was, as expected, absent.

**Figure 3.**
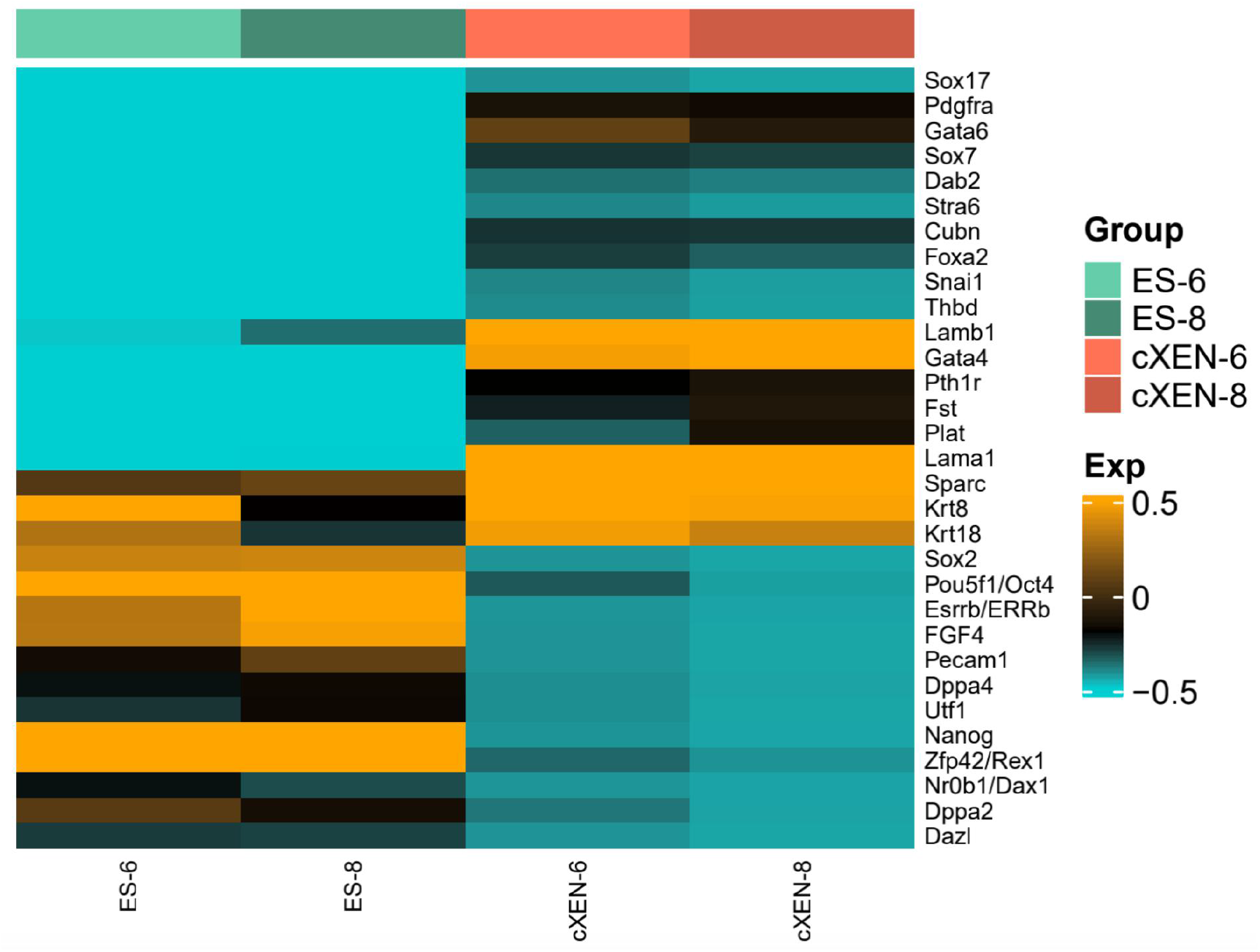
NanoString Gene Expression Analysis. (A) Heatmap nanoString analysis of two SOX17-deficient ES cell lines (ES-6 and ES-8) and two SOX17-deficient XEN cell lines (cXEN-6 and cXEN-8). The SOX17-deficient cXEN cell lines are indicated in red. Heatmap colors correspond to log_10_ values of normalized counts as indicated in the color key, from dark turquoise (low) to orange (high).

### SOX17-Deficient XEN Cells Contribute to the Parietal Endoderm

To test their *in vivo* potential, we injected cells of one SOX17-deficient cXEN cell line (cXEN-13), which has a GFP+ reporter (R26-tauGFP41 × SOX17-Cre), into blastocysts of C57BL/6J origin and transferred the injected blastocysts into pseudopregnant recipients. We transferred 38 blastocysts injected with cXEN-13 cells, identified 30 implantation embryos at E6.5-E7.5, and recovered 24 embryos, among which there were 9 chimeras, and we recovered 24 embryos, 9 of which (37.5%) had GFP+ cells contributed to their parietal endoderm (Fig. 4A-C).

**Figure 4.**
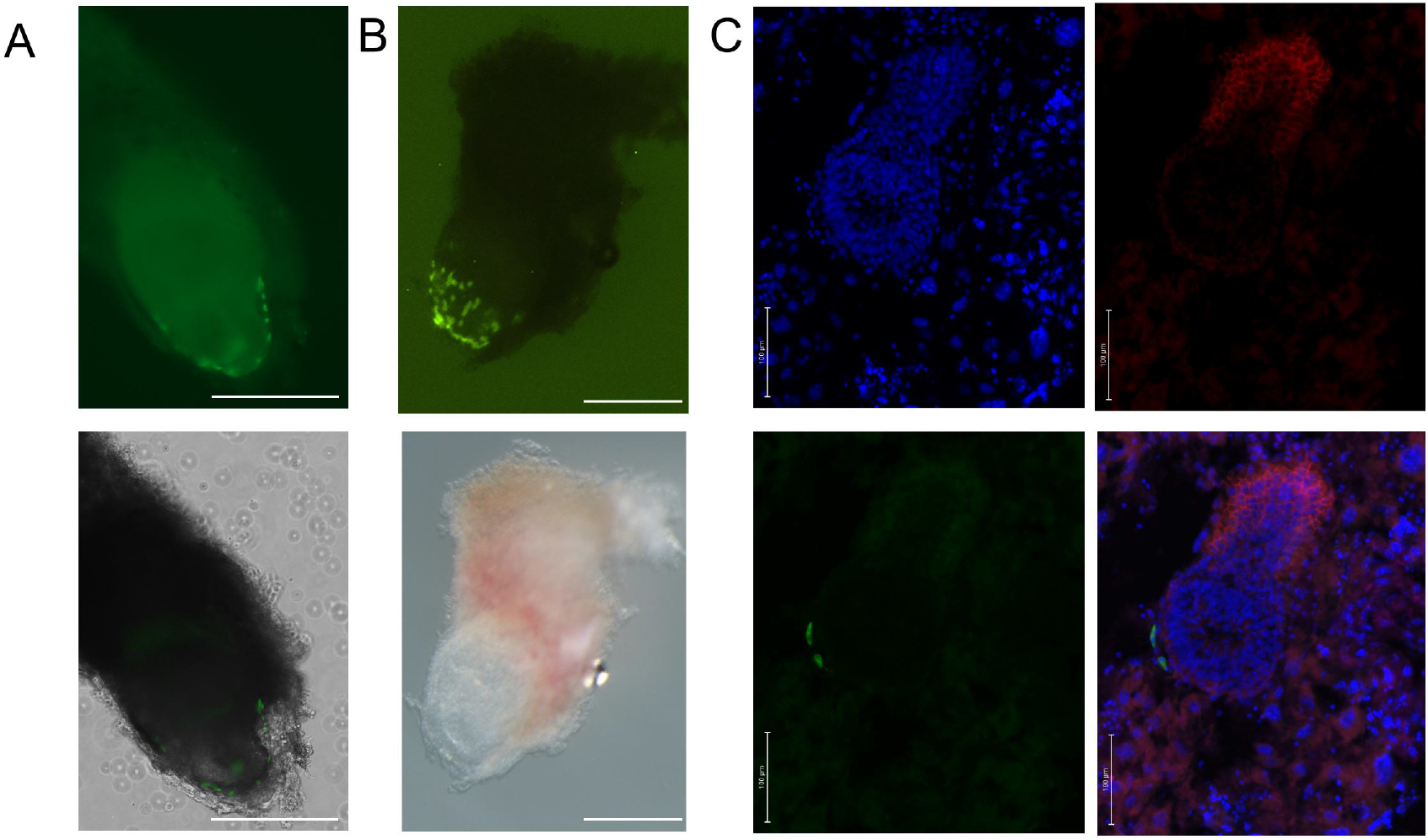
SOX17-Deficient XEN Cells Contribute to the Parietal Endoderm. (A) A wholemount of an E6.5 chimeric embryo was imaged in brightfield and fluorescence, (B) A wholemount of an E7.5 chimeric embryo was imaged in brightfield and fluorescence using a Nikon SMZ18 fluorescence microscope. (C) A section of the decidua of another E7.5 embryo, showing the merged image of fluorescence from DAPI, GFP, and E-cadherin (red), was imaged using a Zeiss LSM 710 (H). Scale bars: 400 *μ*m in A and B panels, 100 *μ*m in C panel.

## DISCUSSION

We derived 6 SOX17-deficient XEN cell lines: two XEN cell lines (X-ICM-12 and X-ICM-16) and four cXEN cell lines (cXEN-13, cXEN-15, cXEN-6 and cXEN-8).

Why are SOX17-deficient XEN cell lines not easy to derive from preimplantation embryos (Niakan et al., 2010)? The missing SOX17 signal reduces the number of PrE cells in blastocysts (Artus et al., 2013). These remaining PrE cells still have the ability to support the SOX17-null concepti to develop to E8.5 and E9.5 (Igarashi et al., 2018). The SOX17-null concepti showed high levels of GATA6 expression in PE cells at E8.5 and E9.5 (Igarashi et al., 2018). This result indicated that the model of sequential expression of PrE lineage-specific genes is Gata6 > Sox17 (Artus et al., 2010, 2011). We observed that GFP+ cells could be maintained in culture and grew slowly to form large colonies. However, in the mixed ES-XEN cultures that we derived from blastocysts, ES cells grew much faster than XEN cells, and ES cells dominated after several passages.

Why were we able to chemically convert SOX17-deficient ES cell lines into cXEN cell lines whereas Cho et al., 2012 were not? First, we used a method called the crowded culture method or extended the culture method (Lin et al., 2017). We changed the medium every other day but passaged the cells every 1-2 weeks. The conventional method is to passage cells frequently (Niakan et al., 2013). We observed that SOX17-deficient ES cells were more difficult to convert than SOX17-Cre heterozygous ES cells in TS cell medium with F4H. Second, we collected cells suspended in the culture medium and spun down the medium to enrich for the XEN-like cells after plating into new dishes. We found that XEN cells cultured in TS cell medium were easier to be collected in suspension than in ES medium when colonies became crowded. The conventional method to change medium and passage cells entails removing the culture medium, which would also remove the suspended (XEN-like) cells.

The model of sequential expression of PrE cell lineage-specific genes is *Gata6* > *Pdgfra* > *Sox17* > *Gata4* > *Sox7* (Artus et al., 2010), which is consistent with the failure in Gata6 mutant embryos to activate sequential expression of Pdgfra, SOX17, and Gata4 in the PrE of blastocysts (Schrode et al., 2014). Gata6 mutants exhibit a complete absence of PrE, while SOX17 or PDGFRA mutants exhibit only a reduced number of PrE cells (Artus et al., 2011, 2013; Schrode et al., 2014). This means that SOX17 or PDGFRA mutants could be partially rescued by other genes or pathways. We speculate that PDGFRA and SOX17 coud have parallel expression, because SOX17- or PDGFRA-deficient cells in blastocysts cannot block each other’s expression (Artus et al., 2011, 2013). We speculate that PDGFRA-deficient XEN cells could be rescued by SOX17 with parallel expression and, conversely, that SOX17 mutant cells could be rescued by PDGFRA in the PrE and in XEN cell derivation and maintenance with parallel expression.

The SOX17-deficient XEN cell lines are healthy, grow as well as wild-type and SOX17-Cre heterozygous XEN cell lines, and differ thus far only in *SOX17* expression from SOX17-Cre heterozygous XEN cell lines. The rate of chimeras among recovered embryos (37.5%) was similar to that obtained with other genetically marked pre- and post-XEN cell lines (35-39%) (Lin et al., 2016). Further experiments, such as RNA-seq, may reveal differences in gene expression between SOX17-deficient and wild-type XEN cell lines.

## EXPERIMENTAL PROCEDURES

### Mouse Strains

The SOX17-Cre strain (Choi et al., 2012) was obtained from MMRRC, strain 036463-UNC, strain name SOX17<tm2(EGFP/cre) Mgn>/Mmnc. Although this strain is reported to contain and express GFP, and we confirmed the presence of GFP in this targeted insertion in the SOX17 locus, but we cannot detect GFP expression in embryos and cell lines derived from them. The R26-tauGFP41 reporter strain (Wen et al., 2011) was obtained from Dr. Uli Boehm, Universität des Saarlandes, Homburg, Germany.

### TS Cell Medium

Advanced RPMI-1640 (Gibco #12633-012) was supplemented with 20% (vol/vol) FBS (HyClone #SH30071.03), 2 mM GlutaMAX Supplement (Gibco #35050), 1% penicillin/streptomycin (Specialty Media #TMS-AB2-C), 0.1 mM β-mercaptoethanol (Gibco #21985-023), and 1 mM sodium pyruvate (Gibco #11360-039) and with F4H, which consisted of 25 ng/ml FGF4 (Peprotech #100-31) and 1 *μ*g/ml heparin (Sigma #H3149).

### ES Cell Medium

DMEM (Specialty Media #SLM-220) was supplemented with 15% FBS (HyClone #SH30071.03), 2 mM GlutaMAX Supplement, 1% penicillin/streptomycin, 1% β-mercaptoethanol (Specialty Media #ES-007-E), 0.1 mM nonessential amino acids (Gibco #11140-035), 1 mM sodium pyruvate, and 1000 IU/ml leukemia inhibitory factor (LIF) (Millipore #ESG1107).

### cXEN Cell Conversion from ES Cells with Retinoic Acid and Activin A

The chemical conversion was performed as described (Cho et al., 2012), with modifications. In the XEN culture medium, we increased fetal bovine serum from 13% to 20% and added 1 mM sodium pyruvate. ES cells were cultured in ES medium with LIF until they reached 70-80% confluency and then in standard TS medium with 25 ng/ml FGF4 and 1 *μ*g/ml heparin (F4H). After 24 h, the medium was changed to TS medium with F4H, 0.01 *μ*M all-trans retinoic acid (Sigma #R2625) and 10 ng/ml Activin A (R&D Systems #338-AC-010). After 48 h, we changed the culture medium to TS medium with F4H. Cells were maintained hereafter in standard TS medium with F4H. After 24 h, cells were dissociated with TrypLE Express and plated at a 1:2 dilution in a dish coated with gelatin and with or without MEF. On approximately day 15, a fraction of XEN-like cells did not adhere tightly to the dishes. We collected the culture medium into a 2.0 ml Eppendorf tube, centrifuged the tube for 30 sec in a minicentrifuge, and removed the supernatant. We washed the dishes twice with calcium- and magnesium-free PBS, transferred the PBS with suspension cells to the tube, centrifuged the tube, and removed the supernatant. Finally, we added fresh medium to the tube and transferred the medium, including the pelleted cells, back into new dishes coated with gelatin and covered with MEF. We applied this method to collect XEN-like cells every day while changing the medium.

### Immunofluorescence and Imaging

Cell lines were cultured in 24-well dishes. Cells were fixed in 4% paraformaldehyde at room temperature for 30 min, permeabilized with 0.1% Triton X-100 in PBS (PBST) for 30 min and blocked with 5% normal donkey serum (Jackson ImmunoResearch Laboratories, #017-000-121) diluted in PBST (blocking solution) for 1 hr. Primary antibodies were diluted at 1:200 in blocking solution, and samples were incubated at 4°C rotating overnight. After three 10-min washes in PBST, samples were incubated for 1-1.5 hr at room temperature in a 1:500 dilution of secondary antibody in blocking solution, washed and covered with PBST containing DAPI. Primary antibodies from Santa Cruz Biotechnology were against GATA4 (#SC-1237), DAB2 (#SC-13982), OCT3-4 (#SC-5279), NANOG (#SC-376915), and CDX2 (#SC-166830). Primary antibodies from R&D Systems were against GATA6 (#AF1700), SOX7 (#AF2766), SOX17 (#AF1924), and PDGFRA (#AF1062). Secondary antibodies from Jackson ImmunoResearch Laboratories were Cy5 AffiniPure Donkey antigoat IgG (H+L) (#705-175-147). Secondary antibodies from Invitrogen were donkey anti-rabbit IgG (H+L) with Alexa Fluor 546 (#A10040) and donkey anti-mouse IgG with Alexa Fluor 546 (#A10036).

### NanoString Multiplex Gene Expression Analysis

Cells were collected by trypsinization and centrifugation. Cell pellets were dispensed in RNAlater Stabilization Solution (Qiagen) and stored at −80°C for later use. Cell pellets were lysed in RLT Lysis Plus Buffer using a TissueLyser LT (Qiagen) at 40 Hz for 2 min. Extraction of total RNA was performed using the RNeasy Plus Micro kit (Qiagen). The custom nanoString CodeSet “Extra” was used; sequences of relevant capture and reporter probes are in the Supplementary Information. An aliquot of 100 ng was hybridized at 65°C for 18 hr and processed with nCounter (nanoString Technologies). Background subtraction was performed using the maximum count of the negative control. A two-step normalization was performed: (1) the geometric mean of positive controls was used as the normalization factor across samples, and (2) the geometric mean of *Actb* and *Gapdh* counts was used as the biological reference normalization factor. A heatmap was generated using the heatmap.2 function in the R package gplots.

## AUTHOR CONTRIBUTIONS

J.L. and X.D. designed research, J.L. and A.D performed experiments. J.L. and X.D. analyzed data and wrote the manuscript.

## ACKNOWLEDGMENTS

The study was supported by grants from Senior Talents Project of Yunnan Health Commission (L-2018006), Academician Expert Workstation of Yunnan Province (202005AF150033), Clinical Research Center for Gynecological and Obstetric Disease of Yunnan Province (2022YJZX-FC21) and the National Natural Science Foundation of China (31970823).

